# The Unconventional Self-Cleavage of Selenoprotein K

**DOI:** 10.1101/2021.05.15.444318

**Authors:** Rujin Cheng, Jun Liu, Martin B. Forstner, George Woodward, Elmer Heppard, Peter R. Hoffmann, Sharon Rozovsky

## Abstract

Through known association with other proteins, human selenoprotein K (selenok) is currently implicated in the palmitoylation of proteins, degradation of misfolded proteins, innate immune response, and the life cycle of SARS-CoV-2 virus. However, neither the catalytic function of selenok’s selenocysteine (Sec), which, curiously, resides in an intrinsically disordered protein segment nor selenok’s specific role in these pathways are known to date. This report casts these questions in a new light as it describes that selenok is able -both *in vitro* and *in vivo-* to cleave some of its own peptide bonds. The cleavages not only release selenok segments that contain its reactive Sec, but as the specific cleavage sites were identified, they proved to cluster tightly near sites through which selenok interacts with protein partners. Furthermore, it is shown that selenok’s cleavage activity is neither restricted to itself nor promiscuous but selectively extends to at least one of its protein partners. Together, selenok’s cleavage ability and its features have all hallmarks of a regulatory mechanism that could play a central role in selenok’s associations with other proteins and its cellular functions overall.

## Introduction

Selenok is a small intrinsically disordered protein (IDP) in the family of selenoproteins, whose members all contain the genetically encoded amino acid selenocysteine (Sec, U). Sec is highly reactive, and all well-characterized selenoproteins are enzymes with this amino acid in a central role of their mechanisms (Arnér, 2010). Many selenoproteins are involved in the management of reactive oxygen species (ROS), where Sec’s reactivity is exploited to tame these signaling molecules that -if left unchecked-can cause severe cellular damage. However, harvesting Sec’s reactive benefits puts a high burden on cellular resources. Its production and insertion require tight control and utilize specialized biosynthetic machinery, without which selenocompounds would aggressively react with their environment and soon inflict irreversible damage to cellular components (Hatfield et al., 2014).

Selenok’s cellular and biochemical functions are largely undetermined, and its enzymatic activity unknown (Liu & Rozovsky, 2015). Although we have previously shown that on its own, it does not have efficient oxidoreductase activity (Liu et al., 2014), given selenoproteins’ track record and the cellular cost of Sec insertion, selenok is most likely also an enzyme. Curiously, this would put selenok in a very exclusive group, as only a handful of intrinsically disordered proteins are known to possess enzymatic activity (Schulenburg & Hilvert, 2013). To develop an idea of selenok’s function, one should consider selenok’s known associations and interactions with other proteins. Through these, selenok contributes to protein quality control, protein palmitoylation, immune response, and possibly partners with the 3a protein of the SARS-CoV-2 virus (Gordon et al., 2020; Liu & Rozovsky, 2015; Stukalov et al., 2021).

In the palmitoylation pathway, which adds fatty acids to proteins, selenok binds and modulates its protein partner palmitoyltransferase DHHC6 (Fig. 1A left panel) through interactions that depend on Sec (Fredericks et al., 2017; Fredericks et al., 2014). It was shown that interrupting this interaction result in impaired cellular Ca^2+^ signaling. Its role in the palmitoylation pathway is the proposed reason for selenok’s demonstrated involvement in cancer metastasis, turning it into a target of new treatment strategies for prostate cancer (Liu & Rozovsky, 2015; Marciel & Hoffmann, 2019; Pitts & Hoffmann, 2018).

**Figure 1.**
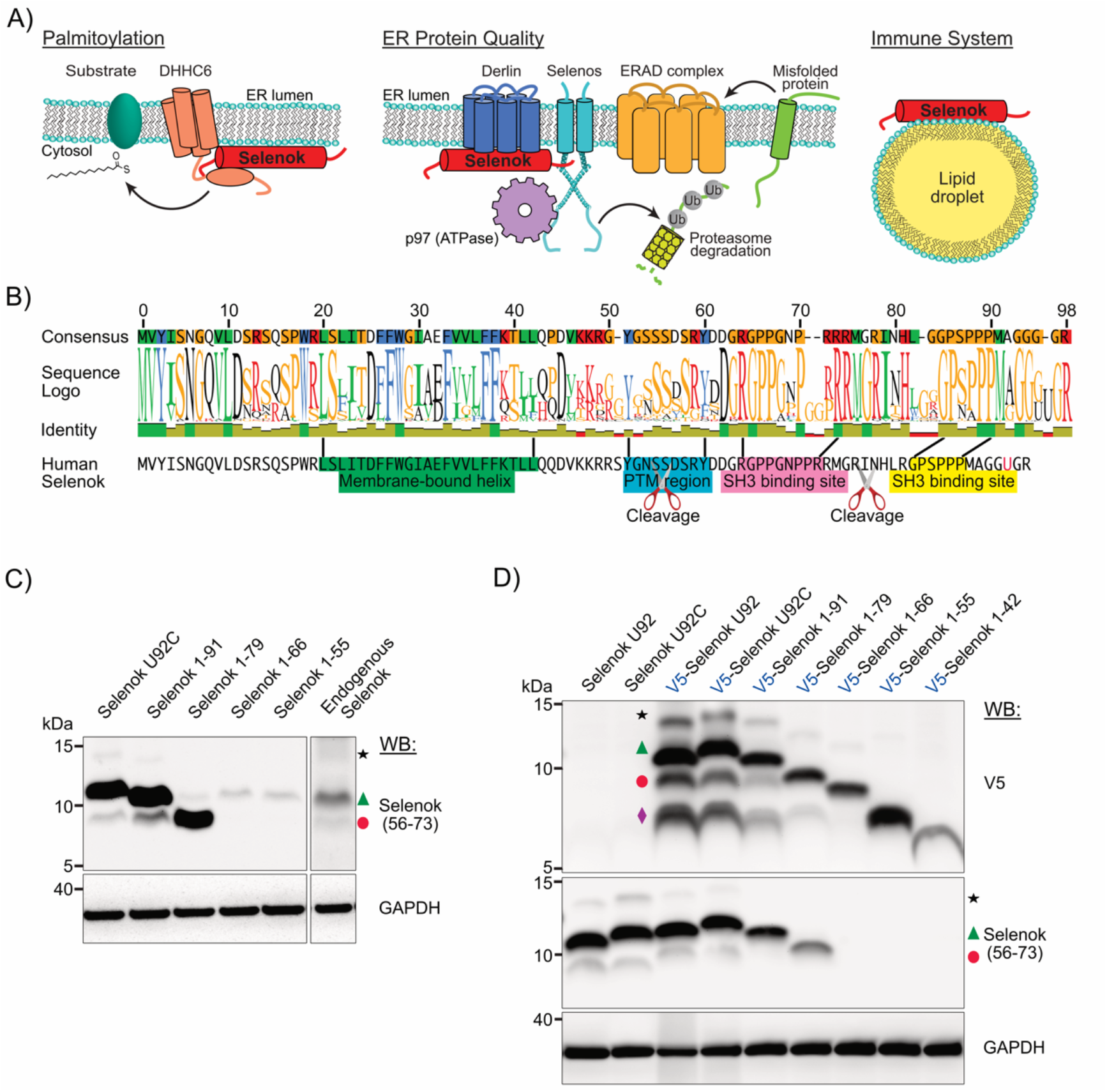
Shorter forms of selenok in HEK 293T. A) Known cellular roles of selenok (red): Involved in protein palmitoylation (left), member of a multiprotein complex at the ER membrane translocating misfolded ER proteins for degradation (middle), and contributing to the innate immune response when associated with lipid droplets (right). B) Logo representation of the sequence alignment of 50 mammalian selenoks. Among the many strongly conserved regions of selenok, its membrane-bound helix (dark green) is the most conserved segment (see also Fig. S1). Bottom line: the sequence of human selenok. C) Selenok detection in HEK 293T lysates in western blots using the anti-selenok (56-73) antibody. Right lanes: over-expressed tag-less selenokof different lengths serve as reliable molecular weight markers. Left lane: endogenous selenok. The resulting 3 major bands are identified as: the full-length selenok detected in all samples (green triangle), a shorter selenok form (red circle), and full-length selenok with an N-linked glycosylation at Asn54 (black star). D) Detection of tag-less and N-terminally V5-tagged selenok over-expressed in HEK 293T cells by anti-selenok (56-73) and anti-V5 antibody western blots. Over-expressed V5-selenok variants of different lengths served as reliable molecular weight markers. V5-selenok samples contain from top to bottom: glycosylated full-length selenok (black star), full-length selenok (green triangle), a short form of selenok running similar to V5-selenok 1-79 (red circle), and a shorter form similar in weight to the V5-selenok 1-55 variant (purple diamond). When overexpressed, selenok U92 runs lower than selenok U92C (left two lanes) because limitations of cellular Sec incorporation lead to frequent truncation at position 92, resulting in selenok 1-91.

Selenok is also a member of the endoplasmic reticulum-associated degradation (ERAD) pathway (8-10), which extracts defect proteins and protein complexes from the ER’s membrane and lumen, transports them the cytoplasm, and marks them via polyubiquitylation for their final degradation by the proteasome (Fig. 1A middle panel). In the ERAD, selenok interacts with derlin and a complex formed by the ATPase p97 and selenoprotein s (selenos) (Shchedrina et al., 2011). While selenok’s role in the ERAD pathway remains to be resolved, it appears to be important enough that silencing of the selenoprotein impairs the function of the pathway and results in the accumulation of dysfunctional proteins not properly processed (Lee et al., 2014). Through its ERAD association, selenok is also coupled to the cellular response to oxidative stress, and as such stress increases, cellular levels of selenok do indeed rise (Du et al., 2010; Touat-Hamici et al., 2014).

With lipid droplets, a third cellular location of selenok has been recently experimentally recognized (Fig. 1A right panel). These independent organelles have a core of neutral lipids that is coated by a monolayer of lipids and proteins. Such lipid droplets not only act as storage of metabolic energy in the form of lipids, but they also coordinate multiple reactions such as lipid biosynthesis and signaling (Gao & Goodman, 2015; Thiam et al., 2013). Selenok’s function in the droplets is also unknown, but it appears to be connected to the innate immune response, as, upon its activation, selenok in lipid droplets is upregulated (Bosch et al., 2020).

Most likely, selenok plays additional, as of yet undiscovered, roles in the cell as the human protein atlas, for example, places selenok also in the nucleoplasm. Additional hints regarding selenok’s functions can come from selenok’s sequence, which contains three distinct regions: A short N-terminal segment (residues 1-19), a hydrophobic helix (residues 20-42), and an intrinsically disordered C-terminal segment (residues 43-94). Of these, the hydrophobic segment is the most evolutionary conserved one (Fig. 1B and Fig. S1), which originated selenok’s initial identification as a single-pass membrane protein. However, the helical projection of this segment strongly suggests that it is most likely an amphipathic helix (Fig. S2), designating selenok as a peripheral membrane protein that is bound to one lipid leaflet but does not cross the membrane. Selenok’s sequence is rich in molecular recognition features (MoRFs, also referred to as short linear motifs (SLiMs)) (Van Roey et al., 2014). A frequent feature of IDPs, these motifs dictate their cellular behavior. Using the Eukaryotic Linear Motif (ELM) resource (Kumar et al., 2020), the following motifs can be established with high probability: two Src-homology 3 (SH3) binding sequences, of which one was shown experimentally to interact with DHHC6 (Fredericks & Hoffmann, 2015), two Src-homology 2 (SH2) and a 14-3-3 recruiting sequences. Thus, selenok has the potential to bind and interact with many proteins, only a few of which have been identified at this point.

Regardless of selenok’s specific role, this work puts selenok’s interactions with other proteins into a new light. It will not only be shown that selenok can cleave itself, that the preferred cleavage sites are at strategic locations that would selectively interrupt protein interactions, that such self-cleavage always liberates the C-terminal Sec, thus terminating its local action, but also that selenok can selectively cleave a protein partner. These newly identified features imbue selenok with active ways to potentially control interactions with and functions of other proteins.

## Results

### Shorter forms of selenok are present in HEK 293T

When fragments of small intrinsically disordered proteins, such as selenok, are observed in cells, they are typically considered an insignificant byproduct of IDPs’ susceptibility to proteolytic cleavage. Yet, when working with selenok, the consistencies regarding its fragments were quite notable and suggested that selenok might exists in several distinct forms in the cell. To understand if selenok is indeed cleaved to shorter forms in its native environment, where it is surrounded by its protein partners and associated complexes, we choose to take a systematic look at selenok in HEK 293T cells. Not only is HEK 293T a well-established model cell line with a track record of selenok studies (Lee et al., 2015; Shchedrina et al., 2011), but it has also been shown that in these cells, calpain II does not cleave selenok (Huang et al., 2011), which eliminates one cause of fragmentation from the start. To detect selenok, a commercial antibody raised against an 18 amino acid peptide of selenok residues 56-73 was used. Because of the possibility of cleavage in the epitope region, we first validated anti-selenok (56-73) detection using the shorter variants 1-42, 1-55, 1-66, 1-79, and 1-91 overexpressed in HEK 293T (Fig. 1C, Fig. 1D, and Fig. S1). In another series of experiments, an N-terminal V5 affinity tag was added to these short selenok variants to independently test for the proteins’ successful expression and exploit them as molecular weight markers in other experiments (Fig. 1D). From such blots in Figure 1C and D (see also Fig. S3), it is straightforward to conclude that a selenok form must contain at least residues 66-73 to be effectively detected by the antibody.

In a western blot of HEK 293T cell lysate containing only endogenous selenok, this antibody labeled three distinct bands (Fig. 1C). To determine the molecular weights of the endogenous selenok in these bands, we exploited overexpressed full-length selenok and the above-mentioned shorter selenok variants as molecular weight markers (for a discussion of the necessity of this approach, please see supporting information). The top band (black star) in Figure 1C is full-length selenok glycosylated on residue N54 (Fig. S4), the band around 10.5 kDa corresponds to full-length selenok (green triangle), while the lower band around 9 kDa is at a similar molecular weight as the variant selenok 1-79 (red circle). Thus, at least one shorter form of endogenous selenok is present in HEK 293T.

Studies of endogenous selenok that freely interacts with native protein partners and complexes at normal expression levels would certainly provide the best insight into its cellular behavior. Unfortunately, full-length selenok is a low abundance protein. Therefore, the even smaller amounts of any derived forms quickly push their characterization in HEK 293T cells beyond current technical abilities. In addition, the only high-binding affinity antibody is the anti-selenok (56-73) that is unable to effectively bind any variants shorter than selenok 1-66, as established above. Thus, to expand the range of detectable selenok variants, we overexpressed selenok with an N-terminal V5 tag. This affinity tag is less charged and contains fewer hydrophobic residues than other commonly used tags and, thus, its use minimizes interactions with membranes or selenok itself. A tag-less selenok was also used to validate that the presence of the V5 tag did not change the overall pattern of selenok variants as detected by the anti selenok (56-73) antibody. Because of the low throughput of the cellular selenocysteine incorporation machinery, the majority of selenok U92 is terminated at position 92 when overexpressed. Thus, we examined not only the native selenok U92 but also its cysteine variant selenok U92C. When selenok with an N-terminal V5 tag that was overexpressed in HEK 293T is detected with an anti-V5 antibody, three major bands appear, as seen in Figure 1D (see also Fig. 3C). The bands correspond to full-length selenok U92C and two shorter selenok forms. Again, using overexpressed V5-selenok variants of different defined lengths as markers, it could be established that the molecular weights of the observed shorter forms are close to selenok 1-79 and 1-55, respectively.

Our findings establish that in HEK 293T selenok is cleaved at least at two major sites. Shorter forms detected by their N-terminal V5 tags were of similar molecular weight as the 1-79 and 1-55 variants. Because they retained their affinity tag, cleavage took place in the C-terminal segment past the amphipathic helix. The fact that the cleavage patterns for endogenous selenok, overexpressed native, and V5-tagged selenok showed little deviation (Fig. 1C-D and Fig. S3) makes this cleavage and their location a property of selenok endogenous to HEK 293T. The origin of the cleavages, however, remained unclassified, and a systematic approach that can also address the potential involvement of mammalian proteases necessitated more controlled experiments using purified selenok.

### Cleavage is intrinsic to selenok

For the study of the basic properties of human selenok, its expression in E. coli was favored for several reasons. In the bacterium, selenok lacks not only its protein partners and mammalian membrane environment but also the presence of mammalian proteases, thus minimizing interactions that can potentially modify the protein. In addition, E. coli is unable to introduce post-translational modifications, thus eliminating any influence of these chemical modifications on cleavage.

In every instance of selenok U92C expression in E. coli, *in vivo* cleavage and its progression over time were observed (Figs. S3 and S5, see also Fig. 3C). This basic observation was not altered by the absence or presence of small affinity tags nor the protein terminal they were attached to. Additionally, fusion to proteins did not arrest selenok cleavage. Like in HEK 293T, when expressed in E. coli, selenok U92C is present in at least two shorter forms in addition to the full-length protein (Figs. S3 and S5). The molecular weights of the two shorter selenok forms were determined to be close to selenok 1-79 and selenok 1-55, using shorter selenok variants expressed in E. coli for direct comparison.

To see if the robustly observed *in-vivo* cleavage of selenok might be, in fact, an intrinsic property, we proceeded by studying purified protein. In ideal experiments, one would monitor the cleavage behavior of full-length selenok over time. However, cleavage always progressed during the growth of bacteria and standard protein purification schemes and lead to a mixture of selenok forms and difficulties in following the time course of cleavage and reliably interpreting the results. Thus, in order to obtain a more controlled starting point, we developed a preparation strategy where different affinity tags are placed at the N and C-terminus to ensure that only intact protein survives purification. Specifically, we chose to add the *Saccharomyces cerevisiae* VMA intein to the C-terminal of a strep-selenok U92C because intein can be used for purification but also induced to self-excise from selenok without leaving a trace (Fig. 2A) (Batjargal et al., 2015; Debelouchina & Muir, 2017). After two affinity chromatography steps to capture the N-terminal strep and the C-terminal VMA intein, only the full-length selenok U92C was retained. After intein release, strep-selenok U92C was successfully formed as confirmed by intact mass spectrometry (Fig. 2C and 2D). When such strep-selenok U92C was incubated over time and monitored at 25 °C (Fig. 2B), we observed in Tricine-SDS-PAGE that the band of the full-length selenok disappeared over time while the intensity of the smaller forms increased. Furthermore, when strep-selenok U92C was concentrated, cleavage occurred faster (Fig. 2B, see also the later discussion regarding concentrating selenok). To map the sites of cleavage, aliquots from the incubation were assayed over time using intact protein mass spectrometry (Fig. 2C and 2D). We reproducibly detected, in multiple repetitions for different constructs and purification strategies, cleavages at five sites clustered near residue 79: R73/M74, G75/R76, N78/H79, H79/L80, and L80/R81 (see Fig. 2D for a typical deconvoluted mass spectrum, and the one specifically acquired for the sample used to generate Fig. 2B. See also Figs. S7 and S8 for additional examples). Among the five sites in the cluster, cleavages at N78/H79 and H79/L80 are the most common. In addition, a cleavage site between S55/S56 was repeatedly observed (Fig. S6). In contrast to the cluster around residue 79, the S55/S56 location is limited to a single cleavage site, suggesting that it is more stringent.

**Figure 2.**
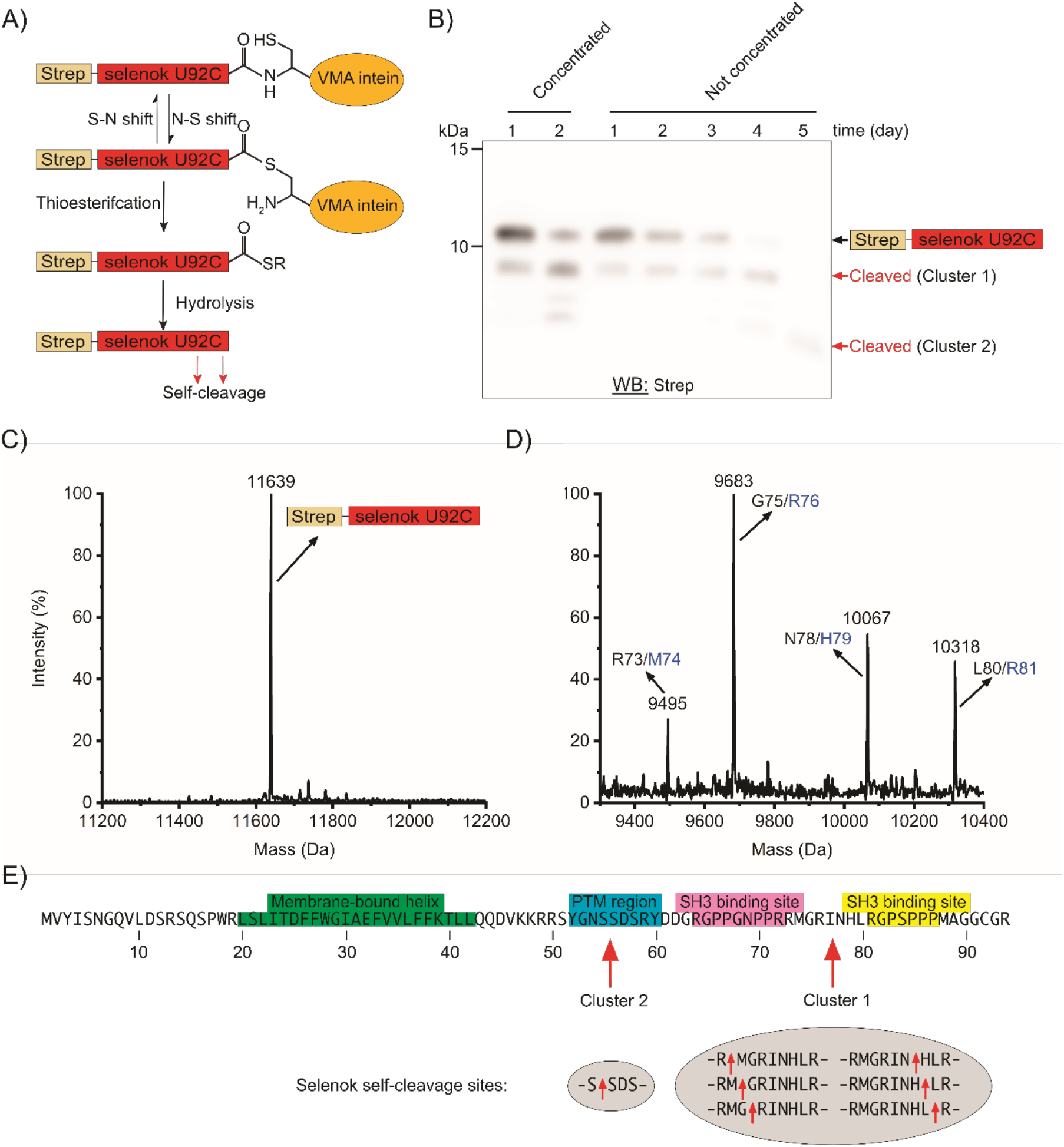
Cleavage of selenok U92C *in vitro* following purification. A) Schematic of full-length strep-selenok U92C preparation from strep-selenok U92C-VMA-intein through sequential StrepTactin and chitin affinity chromatography, followed by intein cleavage and thioester hydrolysis. B) Self-cleavage of purified, full-length strep-selenok U92C overtime at 25°C, detected with a western blot using an anti-strep antibody. Self-cleavage of strep-selenok U92C following concentration (two left lanes) is faster than that of strep-selenok U92C without concentrating (right lanes). C) Deconvoluted ESI-MS spectrum of the strep-selenok U92C following purification. Observed mass 11639 Da corresponds to full-length strep-selenok U92C (calculated 11639 Da). D) Deconvoluted ESI-MS spectrum of strep-selenok U92C following incubation. Detected masses of 9495 Da, 9683 Da, 10067 Da, and 10318 Da correspond to cleavages between residues 73/74, 75/76, 78/79, 80/81, respectively (with a calculated molecular weight of 9495 Da, 9683 Da, 10067 Da, and 10318 Da). E) Location of cleavage sites that were persistently identified using different expression and purification strategies and their position in relation to selenok segments involved in protein interactions.

To assure that the observed cleavage was independent of the purification procedure, maltose-binding protein (MBP) was fused to strep-selenok U92C VMA to prepare strep-selenok U92C using an alternative affinity purification procedure (Figs. S6 and S7). The resulting strep-selenok and its progressive cleavage were identified by intact protein mass spectrometry, as shown in Fig. S7. These results using MBP agreed well with the observations depicted in Fig. 2. In addition, we also purified selenok with small affinity tags from isolated E. coli C43(DE3) membranes. Specifically, selenok U92C with hexahistidine on either the N- or the C-terminus and with an N-terminal strep tag were investigated. Because their yield remained too low for intact protein mass spectrometry, figure S8 contains only their respective western blots. Again, all purified proteins contained shorter forms that can be mapped to cleavages near residues 55 and 79 using the short selenok variant markers. Importantly, in all cases, selenok cleavage was present and continued after purification.

Thus, we were able to establish three aspects of selenok cleavage. Firstly, selenok cleavage occurs even after it has been purified from bacterial cells. Secondly, the overall pattern and rate of selenok cleavage are robust features against all tested variations in the expression and purification schemes to obtain full-length selenok. Finally, there are two main cleavage areas, a single site at residue 55 and a cluster of sites in proximity of residue 79. This overall persistence of the basic observations under the many different conditions we tested does not only render consistent protease contamination highly unlikely but strongly indicates that selenok is, in fact, cleaved by itself.

Interestingly, the cleavage sites at residues 79 and 55 are in segments of selenok that interact with other proteins (red arrows in Fig. 2E). The cluster near residue 79 is between two SH3 binding elements, one of which is used by DHHC6 to interact with selenok (Fredericks et al., 2014). Residue 55 is near predicted binding sites for SH2, SH3, and 14-3-3 proteins (Fig. 3A). Thus, the observed cleavages release these interaction sites, and in a cellular context, such release would terminate selenok interactions with proteins associated with these binding sites. In addition, the segment between residues 52-60 is rich in phosphorylation sites (Hornbeck et al., 2015), further hinting at a possible role of selenok cleavage in cellular signaling. It should also be kept in mind that both cleavages release peptides containing Sec, thus interrupting selenok’s Sec-related enzymatic activity. Furthermore, released segments could potentially move to a different cellular location and act there.

**Figure 3.**
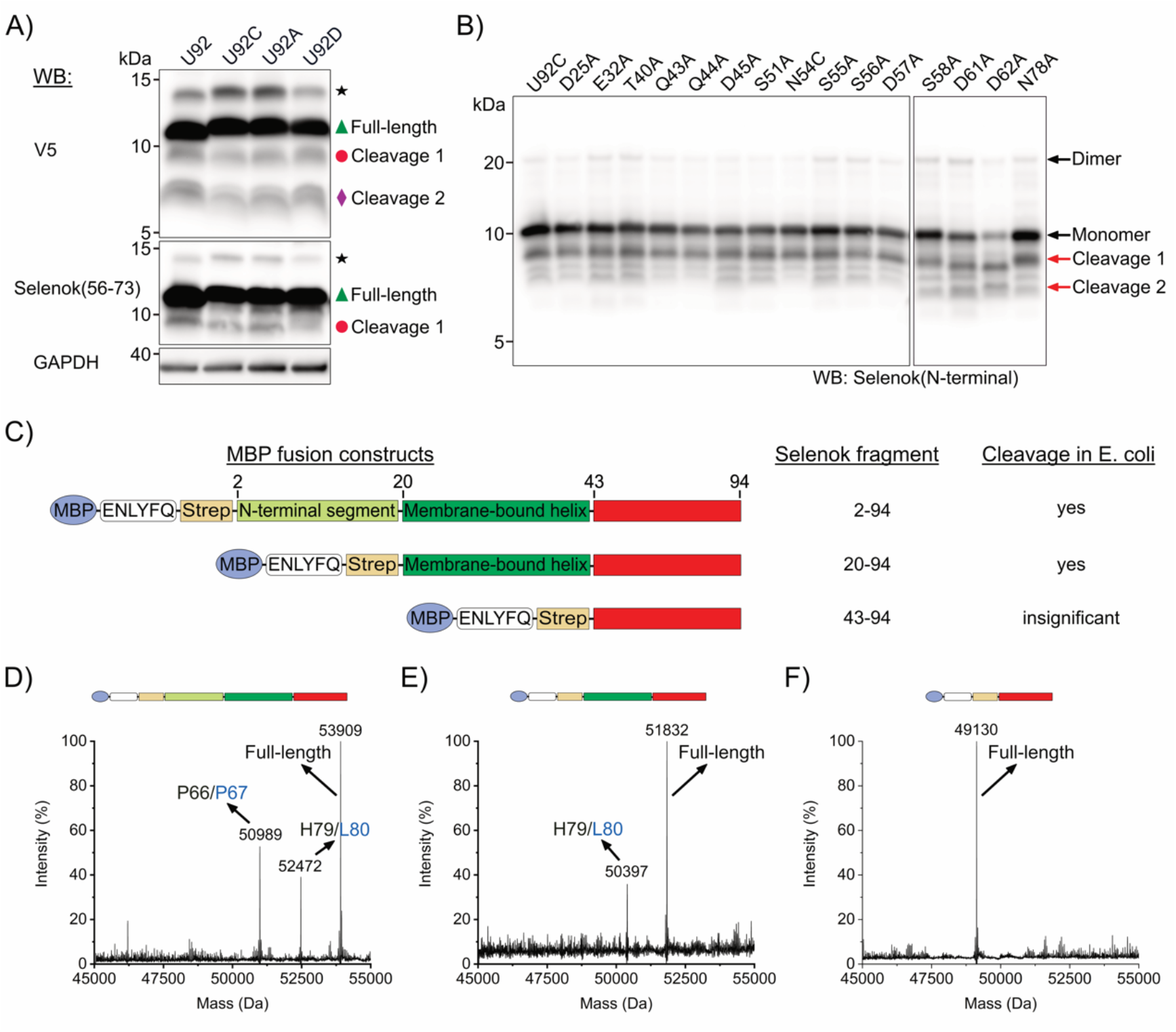
Changes in sequence alter selenok self-cleavage. A) A western blot of V5-selenok with a Sec 92 change to Cys, Ala, or Asp over-expressed in HEK 293T. The bands are: full-length selenok U92C or selenok 1-91 (green triangle), selenok cleaved near residue 79 (red circle), and near residue 55 (purple diamond). The black star indicates glycosylated selenok. B) Western blot of tag-less selenok U92C and point-mutated variants, 20 hours after induction at 18°C in E. coli C43(DE3). The impact of mutations on detection was minimized by employing a customized anti-selenok antibody that binds the unaltered N-terminal region. D) Schematics of selenok variants with different segments and their resulting cleavage activity in E. coli. The necessity of the amphipathic helix for cleavage is evident in the respective deconvoluted ESI-MS right after purification. While for MBP-strep selenok U92C residues 1-94 (D) and MBP-strep selenok U92C residues 20-94 (E) shorter selenok forms are detectable, in MBP-strep selenok U92C residues 43-94 (F) only the full-length protein is present.

### Homing in on selenok’s self-cleavage mechanism

The fact that selenok can cleave itself is particularly intriguing because the selenoprotein is an intrinsically disordered protein (IDP). At this point, only a handful of IDPs are known to be enzymes, and none have shown proteolytic activity (Schulenburg & Hilvert, 2013). However, deducing residues that might be involved in selenok’s cleavage ability from its sequence is hampered by the fact that within selenok’s already tightly conserved sequence, most chemically reactive residues are particularly strictly conserved across species (Fig. 1B and Fig. S1). In addition, selenok does not share homologous structures or sequences with other proteases in the MEROPS database (Rawlings et al., 2018) that may point to a common mechanism. Thus, we set out to further investigate the cleavage to gather further hints regarding the underlying mechanism of cleavage.

To start, we tested whether self-cleavage is unique to human selenok or observed in other mammalian selenok as well. Selenok of *Mus musculus* was chosen because residues close to the cleavage sites (residues 54 and 78) differ from their respective counterparts in human selenok. However, despite these differences, *Mus musculus* selenok purified from E. coli also self-cleaved at similar sites as human selenok (Fig. S9).

Next, possible involvement of the highly reactive Sec at selenok’s C-terminus in the cleavage process was assessed. Selenok was expressed in HEK 293T with its own Sec-coding elements, i.e., the UGA codon and SECIS element. Nevertheless, the limited availability of cellular Sec through its insertion machinery caused the truncation of over-expressed selenok at Sec’s native position 92, as shown in Fig. 1. The resulting mixture of full-length and truncated selenok 1-91 is hard to interpret because the two variants may cleave at different rates. Thus, we compared this selenok with the overexpression of variants in which Sec92 was substituted by either Cys, to retain similar chemical reactivity, Asp, because it mimics Sec’s size and negative charge at physiological pH, or Ala because it has a similar size but is neutral and chemically inert. As apparent in figure 3B, all of these four selenok variants underwent similar cleavages in HEK 293T, indicating that Sec at position 92 is not necessary for selenok’s cleavage activity. This is in line with the earlier observation presented in Fig. 1, where cleavage is observed for both overexpressed selenok 1-91 and selenok 1-79 in HEK 293T. With strong evidence accumulating that residues below position 80 are responsible for cleavage, more controlled *in vitro* experiments using purified selenok were performed. They verified that neither the deletion of the Sec-containing segment UGR (residues 91-94) nor placing different amino acids at position 92 of selenok substantially changed the cleavage (Fig. S10). Thus, it was demonstrated that Sec is not necessary for cleavage activity.

Narrowing down the search for the relevant segments contributing to selenok’s cleavage, several selenok variants of different lengths, with and without the hydrophobic helix, were expressed and purified from E. coli (Fig. 3C-E). From the preservation of general cleavage patterns, it is straightforward to conclude that when residues 1-19 or 80-94 are absent, selenok continues to self-cleave, rendering these segments not necessary for the process. In contrast, variants lacking the amphiphilic helix (selenok 49-94 U92C) were considerably more stable (Fig. 3 and Fig. S11).

With residues 20-79 left as the segment most likely responsible for the cleavage, we first focused on the nucleophilic residues such as Asp, Glu, and Ser in this region. To decipher the role of these residues in the cleavage process, we carried out an Ala scanning mutagenesis survey of tag-less selenok U92C in E. coli (Fig. 3B). The tag-less selenok was selected to assure that tags would not interfere with the segment’s activity. Another concern was the overlap of the region between residue 20 and 70 with the binding site of the commercial anti-selenok antibody. Mutations in that region could selectively alter the binding affinity of the antibody and result in inconclusive or erroneous measurements. Thus, we employed a custom antibody grown to recognize selenok’s N-terminal segment. This antibody binds to the region spanning residues 1-18, which were not altered in the mutagenesis survey. The test included reactive residues such as Ser, Glu, Asn, Lys, and Thr, which were strategically chosen for their proximity to the two cleavage sites or the amphipathic helix. The survey revealed no significant changes in self-cleavage (Fig. 1B and Fig. S12). While histidine is also common in proteases’ active sites, selenok’s only histidine, His79, was not critical for cleavage as the variant selenok H79A U92C underwent self-cleavage (Fig. S12 and S13). In fact, purified selenok H79A U92C was cleaved before and after Ala79 (Fig. S13). Another residue of particular interest is the aspartic acid at position 62. Even in the unusually well-conserved selenok sequence, it stands out as being strictly conserved across all organisms (Fig. 3A and Fig. S1) and suspected – as is its neighboring Asp61 - to play a mechanistic or structural role. In the survey, the cleavage pattern of their alanine variants selenok D61A U92C and selenok D62A U92C are altered (Fig.3 and Fig S12). Both variants, and in particular selenok D62A U92C, exhibited more cleavage near the 55/56 location and a higher ratio of shorter forms to full-length selenok, possibly indicating an increased cleavage rate. Yet, no single mutation stopped cleavage altogether.

We also observed that selenok 20-94 was cleaved while selenok 43-94, which lacks the amphipathic helix (residues 20-42), underwent little cleavage during the expression and purification or even when highly concentrated (Fig. S11). This strongly suggests that the amphipathic helix plays a part in selenok’s cleavage process. This prompted experiments where selenok was successfully purified from membrane fractions of E. coli (Fig. S14). Thus, it was confirmed that selenok was indeed localized to the inner membrane and experiencing a physiological membrane environment. While the helix as a whole is needed for cleavage, substitutions of individual residues within the helix such as S21D, T24A, D25A, E32A, K39A, and T40A did not result in significant changes in the cleavage patterns during expression in E. coli (Fig. 3 And Fig. S12). Since the environment of selenok and particularly that of the amphipathic helix, could potentially modulate cleavage, the influence of the detergent used for protein extraction on cleavage activity was also tested. Self-cleavage was not exclusive to DDM (n-dodecyl-β-D-maltopyranoside) and took also place in other detergents (Fig. S15). Most notably, cleavage appeared as fast in other detergents under similar conditions.

Based on these experiments, there are several lines of evidence that suggest that a conformational change may be involved in accelerating self-cleavage. Firstly, Figure 2G shows that concentrating purified selenok accelerated the cleavage rate (see also Fig. S16). However, concentrating membrane proteins will not only increase protein concentration and thus possibly aid oligomerization but also has the potential to induce changes in the underlying detergent topology, such as transitions from detergent micelles to tubes or other structures (Ferreon et al., 2009). Because selenok cleaves so rapidly, trying to deconvolve the potential contribution of these two effects with biophysical techniques remained fruitless. Unfortunately, this also thwarted our efforts to reliably assess selenok self-cleavage with more traditional assays such as variations of pH or concentrations. However, a second clue comes from fusing selenok to glutathione transferase (GST), which forms a dimer. In these experiments, selenok’s cleavage in E. coli was accelerated to the extent that it was not possible to isolate any full-length selenok (Fig. S17). Finally, when the localization of selenok in E. coli was assayed, the full-length monomer and shorter selenok forms were found only in the membrane fractions. In contrast, the dimeric forms of selenok (besides His_6_-selenok, whose positively charged N-terminal tag can associate with negatively charged lipid head groups) could be found in both the soluble and the membrane fractions (Fig. S14). This suggests that dimeric selenok is more water-soluble than monomeric selenok. If the hydrophobic residues of one amphipathic helix interact with an opposing amphipathic helix, they could be shielded from water and thus straightforwardly explain the observed increased solubility of the selenok dimer. This would also suggest a conformational change driven by the associations of amphipathic helices.

Overall, the data clearly shows that changes in the protein sequence influenced self-cleavage. This further validates that selenok itself is indeed responsible for its cleavage. We have ruled out the Sec, Glu, His, or a Ser residue are the nucleophilic residues. Instead, mutations at Asp residues Asp61 and Asp62 led to a notable change in selenok’s self-cleavage. The amphipathic helix was necessary for appreciable cleavage, and self-cleavage was accelerated by concentrating the protein only if the helix was present.

### Selenok can cleave other proteins

Selenok’s self-proteolysis naturally raises the question of its ability to cleave other proteins. Thus, we incubated selenok with a selection of proteins to probe if its activity extends to other protein substrates. Neither dilute nor concentrated selenok in DDM could cleave p97, beta-lactoglobulin, BSA, thioredoxin, or lysosome regardless of whether they were folded or unfolded (Fig. S18). However, when incubated with selenos, another member of the ERAD complex, selenok cleaved it at specific locations (Fig. 4). Like selenok, selenos is a membrane-bound selenoprotein. It has a hydrophobic helix, a cytosolic soluble cytoplasmic helix, and an intrinsically disordered segment with a highly accessible surface. While selenos’s cytoplasmic portion (cselenos, residues 51 to 189 with a U188C substitution) was stable when incubated alone, it was cleaved by selenok predominantly at the stable alpha-helix (residues 49-123) as opposed to the disordered segment (residues 122-189). The cleavage sites in selenos were clustered in two main locations around Leu58 and Ala67 in contiguous peptide bonds (Fig. 4C and Fig. S19). These cleavage sites are located just before the segment in selenos that is responsible for the recruitment of the ATPase p97 (Tang et al., 2017). Thus, the assay demonstrated that selenok can cleave other proteins, but its activity is not promiscuous.

**Figure 4.**
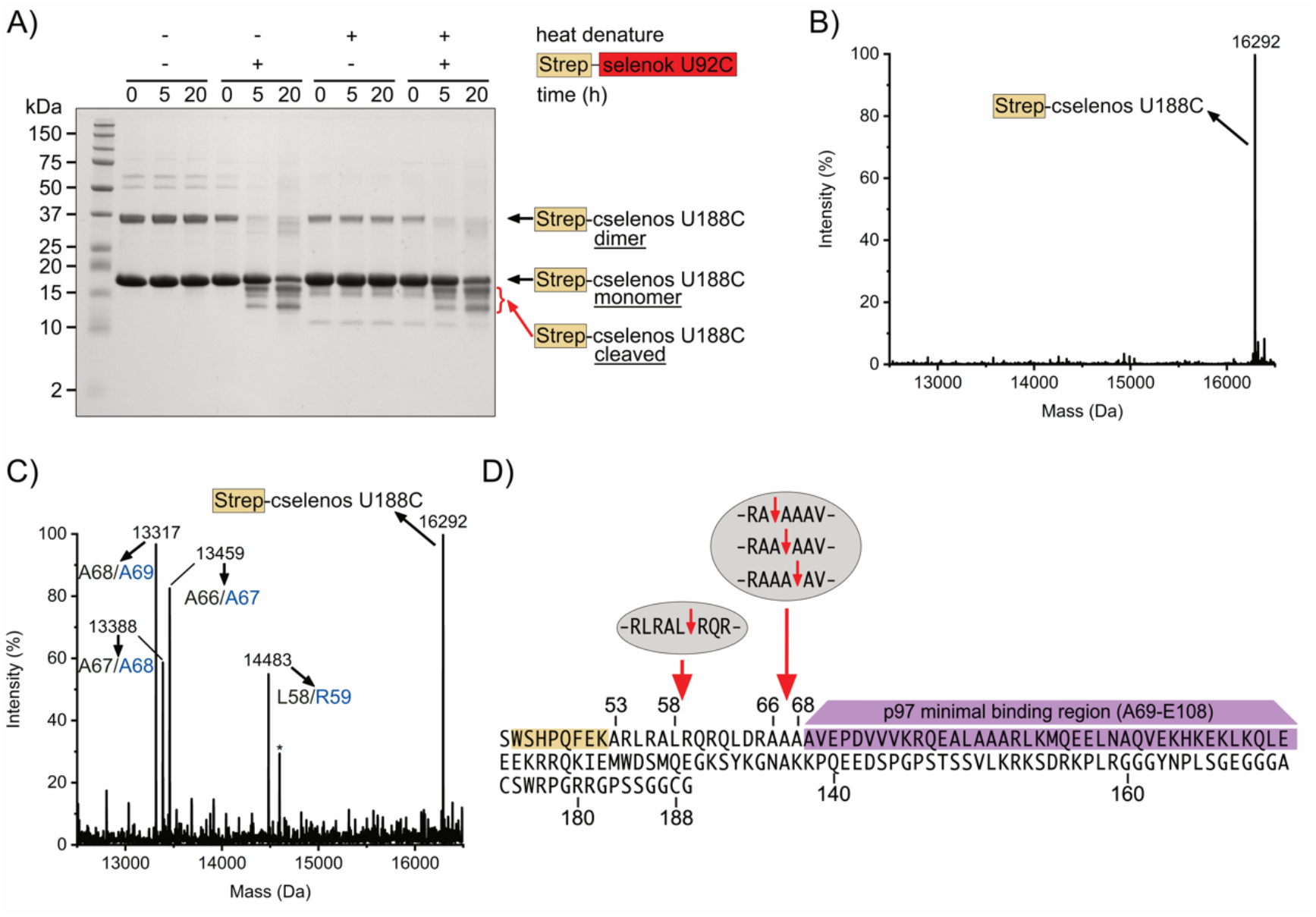
Selenok cleaves its protein partner, selenos. A) Tricine-SDS-PAGE showing incubations at 37°C of heat-denatured and non-denatured strep-cselenos U188C with and without concentrated strep-selenok U92C in DDM micelles. The concentration of strep-selenok U92C is too low to be detected. B) Deconvoluted ESI-MS spectrum of strep-cselenos U188C incubated alone. The molecular mass of 16292 Da corresponds to full-length strep-cselenos U188C (calculated 16293 Da). No shorter forms are detected. C) Deconvoluted ESI-MS spectrum of shorter strep-cselenos U188C variants are generated only when incubating with strep-selenok U92C. Detected masses of 13317 Da, 13388 Da, 13459 Da, and 14483 Da correspond to cleavages between residues 68/69, 67/68, 66/67, and 58/59, respectively (calculated molecular weights are 13319 Da, 13390 Da, 13461 Da, and 14485 Da, respectively). D) Strep-cselenos U188C is cleaved by strep-selenok U92C at two sites (red arrows) in close vicinity to the ATPase p97 binding sequence (purple). The strep affinity tag is light brown. The grey areas above the sequence show the cleavage sites in selenos resulting from incubations with selenok, as identified in multiple biological samples.

## Discussion

We showed that selenok cleaves itself and at least one other protein with a related function (selenos). Self-cleavage was detected *in vivo* in human and bacterial cells under multiple expression conditions and in experiments involving a large array of purified selenok variants regardless of the purification strategy. These results confidently rule out that selenok’s cleavage is caused by an external protease or a specific assay component because it is unlikely that an accidental protease could be consistently present in all these exhaustive purifications. Also ruling an external protease are assays in which many other proteins remained stable when incubated together with selenok. Furthermore, an alanine scanning mutagenesis survey confirmed that changes in selenok amino acid sequence can lead to altered cleavage behavior.

The sites of cleavage in selenok were mapped to the peptide bond between residues 55 and 56 and bonds close to residue 79. Both cleavage sites are strategically located near segments that contain interaction sites with other proteins, such as domains binding with SH3, SH2, and WW motifs. Cleavage at either site releases peptide containing selenok’s Sec. The alanine scanning study showed that the cleavage pattern and rate were shifted by several mutations (most notably, Asp61 and Asp62), while no single mutation completely arrested cleavage activity. However, selenok’s amphiphilic helix is required for cleavage to occur at an appreciable rate, and cleavage is accelerated when selenok is concentrated.

Selenok’s self-cleavage is, of course, not an isolated case. In fact, self-cleavage is a common cellular regulatory mechanism that is employed, for example, for controlling enzyme activity, immune activation, and apoptosis (see abundant examples in the MEROPS database (Rawlings et al., 2018)). However, comparing selenok’s self-cleavage with these examples is problematic. Self-cleavage is often enzymatic and carried out by proteolytic proteins (i.e., proteases) that catalyze peptide bond cleavage by either hydrolysis or elimination (as classified in (Rawlings & Bateman, 2021)). The enzymes that employ hydrolysis commonly use nucleophilic residues such as Cys, Glu, and Asp, but other residues and metals have also been found to contribute to those cleavage mechanisms (Klein et al., 2018). This leaves asparagine peptide lyases, currently the only known nonhydrolytic protease whose asparagine exclusively cleaves its own peptide bond (Rawlings et al., 2011). Selenok is different in several ways. Unlike the self-cleaving proteins above, it lacks a stable tertiary structure. Furthermore, selenok’s cleavage sites are not confined to a single peptide bond but also encompass neighboring bonds (Fig. 2E). Rather than resembling a protease’s self-cleavage, selenok shares commonalities with self-cleaving peptides in proteins that employ chemical proteolysis, exemplified by the peptide-induced self-cleavage of tyrosinase (Kampatsikas et al., 2019) or the self-cleavage of antibodies’ hinge and domain-domain interfaces (Cordoba et al., 2005). Chemical proteolysis is a general term for non-enzymatic cleavage that is mediated by neighboring residues in contact with an exposed protein backbone (Raju, 2019). It is typically driven by β-elimination, such as a nucleophilic attack by an aspartate or isoaspartate residue, the deamidation of glutamine and asparagine residues, or the degradation of disulfide bonds, and can also involve oxidation, isomerization, and light or metal-induced damage (Manning et al., 2010; Reubsaet et al., 1998). Like in the case of selenok, such processes generate a cluster of cleavages in neighboring peptide bonds, they depend on the conformational flexibility at the sites of cleavage, and the respective proteins cannot be stabilized by a single mutation (Harris, 2005). Differences, however, are in the kinetics. Selenok’s cleavage occurs rapidly, while chemical proteolysis in the above examples will take weeks at the physiological pH, buffer conditions and temperatures of our studies (Grassi & Cabrele, 2019). Thus, at this point the many similarities suggest that selenok self-cleavage is most likely a form of chemical proteolysis. However, there are yet unknown mechanistic aspects that cause the intriguing difference in selenok’s cleavage. For one, the cleavage can selectively extend to other proteins. Furthermore, at physiological conditions it occurs at an appreciable rate that in turn can be significantly accelerated that could be brought about by conformational change, membrane properties or protein associations.

The highly reproducible observation that concentrating the protein accelerates cleavage could be related to cellular processes by which proteins self-assemble or form clusters. Many instances of such processes exist, for example, the p62 and TRIM5α distribution into sequestosomes during autophagy (Bjørkøy et al., 2005; Carter et al., 2020) or supramolecular antimicrobial peptide assemblies that trap microbes (Simonson et al., 2020). In this context, it is well documented that selenok is enriched in immune cells (Avery & Hoffmann, 2018), and selenok knock out resulted in impaired immune responses (Verma et al., 2011). Accumulation of selenok in lipid droplets (Bersuker et al., 2018) as part of the innate immune response was also recently reported (Bosch et al., 2020). Therefore, selenok or its shorter forms could undergo a similar clustering process and access a concentration-dependent cleavage in the role they are performing in the human immune response.

Because selenok’s sequence is so highly conserved (Fig. S1), and selenok’s cleavage was observed even in endogenous selenok in HEK293, one might speculate about the functional relevance of selenok’s self-cleavage. Since the shorter form of endogenous selenok was detected in HEK293 cells, we know that at least the hydrophobic fragments remained in the cell. This would imply that these fragments stay embedded in membranes and could continue to interact with protein partners, although their specific associations may now differ from the full-length selenok. Because there are many known instances where protein segments shredded by proteases have cellular functions (Kapp et al., 2009), the fate of peptides cleaved from selenok is equally intriguing. Those peptides contain Sec, which in cells is exclusively used for catalysis. Thus, if the cleaved off peptides are not immediately degraded, they would be free to carry enzymatic activity to different locations.

The observation that selenok can selectively cleave its protein partner, selenos, is remarkable as it might implicate selenok in additional functions in the regulation of other proteins. Selenos is not only a member of the ERAD pathway but also a general scaffolding protein that is involved in vesicle trafficking and NFκβ signaling (Capelle et al., 2021; Liu & Rozovsky, 2015; Turanov et al., 2014). Selenok cleaved selenos near the sequence responsible for recruiting the ATPase p97. Such a strategic cleavage would abolish p97 recruitment by selenos and would suggest that under certain cellular conditions, selenok could regulate selenos function.

In general, the elimination of interaction sites via self-cleavage would certainly change selenok’s association with the respective proteins, while the release of the Sec would have a dramatic impact on selenok’s enzymatic function. These processes could be triggered in response to cellular factors, such as changes in oxidative stress or activation of the immune response, leading to cluster formation and accelerated cleavage as suggested by the concentration dependence and importance of the amphipathic helix. Thus, the observed self-cleavage of selenok, utilizing an unconventional form or proteolysis, appears to have all the hallmarks of a regulatory mechanism.

## Materials and Methods

### Expression vectors

*Homo sapiens* SELENOK cDNA (UniProtKB Q9Y6D0) was purchased from the PlasmID Repository and cloned into the pcDNA3.1 vector using KpnI and PmeI cloning sites. The clone contained the downstream untranslated region (3’-UTR), including the native selenok SECIS element, which is necessary for Sec incorporation (Kryukov et al., 2003). The NEB Q5 mutagenesis kit was used for all mutagenesis. For expression in E. coli, the selenok gene was synthesized by DNA2.0 Inc. with codon optimization for E. coli. Sec codon (TGA) was substituted by Cys codon (TGC) in the DNA sequence as a U92C mutation. Selenok U92C gene was then cloned into the pET28a expression vector using BamHI and EcoI cloning sites. Strep, 6xHis, V5, or Flag tags were introduced to the N-terminal or C-terminal of selenok U92C in pET28a by mutagenesis. The same DNA2.0 optimized selenok U92C cDNA was also cloned into the pMAL-c5X vector with an N-terminal MBP tag using sites BamHI and HindIII. A TEV protease cleavage site (ENLYFQG) followed by a strep tag was inserted between the MBP and selenok U92C to assist purification. The sequence SSS was inserted between MBP and ENLYFQG to promote TEV cleavage. This construct was abbreviated as MBP-strep-selenok U92C (residues 2-94) with the first residue deleted, which reflects N-terminal Met processing *in vivo*. Lastly, the gene of MBP-strep-selenok U92C was cloned into the pTYB1 vector using Ndel and SapI cloning sites, generating a fusion protein with a C-terminal Saccharomyces cerevisiae VMA intein (abbreviated as MBP-strep-selenok U92C-VMA). The sequences of all constructs are provided in the supporting information.

### Cell culture and detection of endogenous selenok by western blot

HEK 293T cells (ATCC) were grown in DMEM medium containing 10% fetal bovine serum (FBS) and 1% penicillin streptomycin, supplemented with 100 nM sodium selenite. The cells were trypsinized, washed twice with cold phosphate buffer, and lysed with cold Pierce IP lysis buffer (25 mM Tris-HCl pH 7.4, 150 mM NaCl, 1% NP-40, 1 mM EDTA, 5% glycerol) supplemented with Halt Protease Inhibitor cocktail (ThermoFisher) at 10 μL/mL. Cells were lysed on ice for 30 minutes with periodic mixing and centrifuged at 13,000 g for 10 minutes at 4°C to pellet down the cell debris. The supernatant of the lysate was collected and analyzed by western blot. For Tricine-SDS-PAGE analysis, 90 μL of the supernatant was mixed with 30 μL of 4x tricine loading dye without reducing reagents (Schagger, 2006). 15 μL of each sample was loaded and resolved on a 16% Tricine-SDS-PAGE. For western blotting, transfer was carried overnight to a PVDF membrane at 20 V according to Schägger’s protocol (Schagger, 2006). The transferred membrane was blocked with 5% BSA in TBST (Tris-buffered saline, 0.1% Tween 20) for 1 h, incubated with anti-selenok (56-73) antibody at 1: 15000 dilution (ThermoFisher catalog number PA5-34420, raised in rabbit) for 1 h, washed for 3 times, incubated with horseradish peroxidase (HRP) conjugated IgG at 1: 40000 dilution (ThermoFisher catalog number 31460, raised in goat) for 1 h and washed for 3 times. The western blot was visualized using a chemiluminescent HRP substrate kit (ThermoFisher catalog number 34075). The same samples were also detected by anti-GAPDH (ThermoFisher catalog number MA515738) as the loading control.

### Transfection and detection of over-expressed selenok constructs in HEK 293T by western blot

HEK 293T cells (ATCC) were grown at 37°C under the same condition in standard 6-well plates. Transient transfections were executed on a scale of 5 μl Lipofectamine 3000 and 2.5 μg of selenok mammalian expression vectors for each well using the recommended protocol by the manufacturer (ThermoFisher). DNA and lipofectamine were mixed in the absence of FBS. Transfections were carried out in the presence of 10% FBS, 1% penicillin-streptomycin, and 100 nM sodium selenite for each well. After 20 hours of transfection, the growth medium was removed, cells were trypsinized, washed, lysed, separated, and a sample of the supernatant was taken for Tricine-SDS-PAGE samples. Samples were frozen in liquid N_2_ and stored at -80°C or immediately subject to analysis. Samples were not heated at any step. Western blots were performed according to Schägger’s protocol (Schagger, 2006). Western blots were detected by anti-selenok (56-73) and anti-GAPDH. For constructs with V5 tags, the blots were also detected by anti-V5 (ThermoFisher catalog number R96025).

### Over-expression and detection of selenok U92C constructs in E. coli

Selenok U92C constructs of various tags in pET28a were transformed into E. coli strain C43(DE3) and grown on a scale of 1 L of LB medium at 37°C supplemented with 1 mM MgCl_2_. When the OD reached 0.6, 0.3 mM of IPTG was used to induce protein expression at 18°C. The OD of C43(DE3) culture was measured at different time points after induction. Expression samples were normalized to contain the same number of cells (equal to a volume of 1000 μl and OD of 0.5). Those samples were spun down at 16,000 g for 2 min to remove the medium. Cell pellets were lysed with 90 μl 4 M urea and 30 μl of 4x tricine gel loading dye. The whole-cell lysates were frozen in liquid N_2_ and stored at -80°C or immediately subject to Tricine-SDS-PAGE without heating process. For western blots, the proteins were transferred from the gel onto a PVDF membrane at 20 V overnight following Schägger’s western blot protocol for small proteins (Schagger, 2006). Transferred blots were detected by anti-selenok (56-73) and imaged with the chemiluminescent HRP substrate kit. Similarly, MBP-strep-selenok U92C and the deletion constructs 2-79, 2-66, and 2-55 in pMAL-c5X were transformed into E. coli strain BL21(DE3) followed by the same expression and sample preparation procedure. The expression gel was visualized by Coomassie staining in addition to western blot using anti-selenok (56-73) detection. The same samples were also detected by HRP conjugated StrepTactin antibody at 1:10000 dilution (Bio-Rad catalog number 161038). To allow detection of endogenous selenok, in Fig.1, endogenous selenok samples were twice more concentrated than those with overexpressed tag-less selenok.

### Sucrose gradient ultracentrifugation and protein enrichment from the E. coli membrane

After induction for 20 hours at 18°C, 1 L of C43(DE3) cells transformed with selenok U92C constructs were centrifuged at 5,000 g for 10 min. The pellet was resuspended in 45 ml buffer composed of 50 mM phosphate, 200 mM NaCl and 1 mM EDTA at pH 7.5. After adding PMSF and benzamidine to 1 mM, cells were opened by homogenizer and centrifuged twice for 30 min at 18,000 g to remove inclusions bodies and unbroken cells. 10 ml of the supernatant was applied on top of a sucrose gradient (8 ml of 0.5 M on top and 8 ml of 1.5 M at the bottom) and centrifuged for 18 h at 100,000 g at 4°C. The separated layers were gently collected as 2 ml fractions from top to bottom using a transfer pipette. 90 μl of each fraction were mixed with 30 μl of 4x tris-tricine loading dye. All fractions were analyzed by western blot using anti-selenok (56-73) antibody. The rest of the supernatant was centrifuged by ultracentrifugation at 100,000 g for 1 hour at 4°C without a sucrose gradient. Membranes of E. coli cells located at the bottom were separated. Membrane proteins were extracted in detergent and subject to affinity purification using the IMAC or StrepTactin (IBA) affinity chromatography.

### Recombinant protein expression and purification

MBP-strep-selenok U92C in pMAL-c5X and MBP-strep-selenok U92C-VMA in pTYB1 were transformed into E. coli strain BL21(DE3) and over-expressed on a large scale in TB medium induced with 0.5 mM IPTG at 18°C. Cells were lysed with a Microfluidics microfluidizer with either PMSF and benzamidine or with a commercial protease inhibitor cocktail. The fusion proteins were purified using an amylose column and eluted in 50 mM sodium phosphate, 200 mM NaCl, 1 mM EDTA, 20 mM maltose, 0.067% DDM, pH 7.5. The proteins were subject to cleavage by TEV protease to remove MBP, generating strep-selenok U92C and strep-selenok U92C-VMA. MBP-strep-selenok U92C-VMA and strep-selenok U92C-VMA were subject to a chitin column for specific intein cleavage initiated by adding 75 mM of sodium 2-mercaptoethanesulfonate (MESNA), which facilitates the N→S shift and promotes protein thioesterification. On the column, intein cleavage proceeded at room temperature for 36 h, releasing MBP-strep-selenok U92C thioester or strep-selenok U92C thioester in Buffer B (50 mM MES, 200 mM NaCl, 1 mM EDTA, pH 6.5). The protein thioesters were hydrolyzed at pH 9.0 in the presence of 200 mM 2-mercaptoethanol (βME) for 30 min (Batjargal et al., 2015; Gates et al., 2013). Derived from VMA intein, MBP-strep-selenok U92C and strep-selenok U92C remained full-length protein with minimal cleavage. After buffer exchange to 50 mM sodium phosphate, 200 mM NaCl, 1 mM EDTA, 0.067% DDM, pH 7.5), the full-length recombinant proteins were used for auto-proteolysis assays by mass spectrometry and SDS-PAGE. Sec incorporation was accomplished by adding selenocysteine to defined growth medium as described in ref (Liu et al., 2012).

### Mass spectrometry

Mass spectra of intact proteins were obtained using a Xevo G2-S QTOF on a Waters ACQUITY UPLC Protein BEH C4 reverse-phase column (300 Å, 1.7 μm, 2.1 mm x 50 mm). An acetonitrile gradient from 5%-95% was used with 0.1% formic acid, over a run time of 5 min and a constant flow rate of 0.5 mL/min at 40°C. The spectra were deconvoluted using MaxEnt1.

### Cleavage assays

Strep-selenok U92C was concentrated in 50 mM sodium phosphate, 200 mM NaCl, 1 mM EDTA, 20 mM maltose, 0.067% DDM, pH 7.5 using a 50 kDa cutoff concentrator (Millipore Sigma). Concentrated strep-selenok U92C was incubated with or without other proteins at 37°C. Samples at different time points were analyzed by Tricine-SDS-PAGE and mass spectrometry. (See detailed cleavage assay of other proteins in supplementary materials and methods).

## Acknowledgments

Research reported in this publication was supported by the National Institute of General Medical Sciences (NIGMS) of the National Institutes of Health under award number GM121607 to S.R. We also acknowledge support by MCB 1616178 from the National Science Foundation. Instrumentation was supported by NIH P20GM104316 and P30GM110758. P.R.H acknowledges NIH grant R01AI089999.

## Author Contributions

R.C. and S.R. designed research; R.C., J.L., M.B.F, G.W., and E.H. performed research; P.R.H. contributed new reagents, R.C. and S.R. analyzed data, R.C., M.B.F, and S.R. and wrote the paper.

## Classification

biochemistry and chemical biology.

